# Tau Filaments from Amyotrophic Lateral Sclerosis/Parkinsonism-Dementia Complex (ALS/PDC) adopt the CTE Fold

**DOI:** 10.1101/2023.04.26.538417

**Authors:** Chao Qi, Bert M. Verheijen, Yasumasa Kokubo, Yang Shi, Stephan Tetter, Alexey G. Murzin, Asa Nakahara, Satoru Morimoto, Marc Vermulst, Ryogen Sasaki, Eleonora Aronica, Yoshifumi Hirokawa, Kiyomitsu Oyanagi, Akiyoshi Kakita, Benjamin Ryskeldi-Falcon, Mari Yoshida, Masato Hasegawa, Sjors H.W. Scheres, Michel Goedert

## Abstract

The amyotrophic lateral sclerosis/parkinsonism-dementia complex (ALS/PDC) of the island of Guam and the Kii peninsula of Japan is a fatal neurodegenerative disease of unknown cause that is characterised by the presence of abundant filamentous tau inclusions in brains and spinal cords. Here we used electron cryo-microscopy (cryo-EM) to determine the structures of tau filaments from the cerebral cortex of three cases of ALS/PDC from Guam and eight cases from Kii, as well as from the spinal cord of two of the Guam cases. Tau filaments had the chronic traumatic encephalopathy (CTE) fold, with variable amounts of Type I and Type II filaments. Paired helical tau filaments were also found in two Kii cases. We also identified a novel Type III CTE tau filament, where protofilaments pack against each other in an anti-parallel fashion. ALS/PDC is the third known tauopathy with CTE-type filaments and abundant tau inclusions in cortical layers II/III, the others being CTE and subacute sclerosing panencephalitis. Because these tauopathies are believed to have environmental causes, our findings support the hypothesis that ALS/PDC is caused by exogenous factors.

**SIGNIFICANCE:** A neurodegenerative disease of unknown cause on the island of Guam and the Kii peninsula of Japan has been widely studied, because patients can suffer from the combined symptoms of motor neuron disease, parkinsonism and dementia. Abnormal filamentous inclusions made of tau protein characterise this amyotrophic lateral sclerosis/parkinsonism-dementia complex (ALS/PDC) and their formation closely correlates with neurodegeneration. Here we have used electron cryo-microscopy (cryo-EM) to show that tau filaments from ALS/PDC are identical to those from chronic traumatic encephalopathy (CTE), a disease caused by repetitive head impacts or blast waves. CTE tau filaments are also found in subacute sclerosing panencephalitis, which is a rare consequence of measles infection. ALS/PDC may therefore also be caused by environmental factors.

## INTRODUCTION

Amyotrophic lateral sclerosis/parkinsonism-dementia complex (ALS/PDC or lytico-bodig) is a fatal disease found in the Chamorro population of Guam (1-4), some families on the Kii peninsula of Japan (5,6), and the Auyu and Jakai people of New Guinea (7). Abundant tau inclusions are present in nerve cells in brains and spinal cords (6,8,9) and are enriched in cortical layers II/III (10,11). Tau inclusions are also found in some glial cells (12). They consist of amyloid filaments that are made of all six brain tau isoforms in a hyperphosphorylated state (8,13). More variably, assembled Aβ, α-synuclein and TDP-43 accumulate too (11,14,15).

The cause of ALS/PDC is unknown, but it is not a simple genetic disorder in an island-bound geographic isolate (16-18). Exogenous factors may play a role in disease aetiology and pathogenesis, supported by the finding that migrants from the Philippines can develop ALS/PDC after living on Guam for more than two decades (19). With increased Westernisation, the incidence of ALS/PDC is decreasing (20). High prevalence of a retinopathy, called linear retinal pigment epitheliopathy (LRPE), has been reported in Guam and Kii ALS/PDC (21,22), similar to infestation by a migrating parasite larva. Both disorders have declined in parallel, suggesting a possible link between ALS/PDC and LRPE.

Tau filaments made of all six brain isoforms in a hyperphosphorylated state are also found in Alzheimer’s disease (AD) and in chronic traumatic encephalopathy (CTE) (23,24). They are found predominantly in cortical layers V/VI in AD (25) and in layers II/III in CTE (26). The latter is caused by repetitive head impacts or exposure to blast waves (27). By cryo-EM, we have shown that tau filaments from AD and CTE each consist of two identical C-shaped protofilaments that comprise residues 306-378 (in the numbering of the 441 amino acid tau isoform) (28-30). They differ by the presence of a hydrophobic cavity in the CTE fold, which encloses a non-proteinaceous density of unknown identity that may be involved in giving rise to this fold. Besides AD, the Alzheimer tau fold also characterises primary age-related tauopathy, familial British dementia, familial Danish dementia and some prion protein amyloidoses (31,32). Besides traumatic encephalopathy syndrome, the CTE tau fold is characteristic of subacute sclerosing panencephalitis (SSPE) (33). The latter is a fatal disorder of the central nervous system that is a rare consequence of infection with measles virus and manifests itself after a symptom-free period of several years (34). Tau inclusions in SSPE are also enriched in cortical layers II/III (35). Here, we report that the CTE fold is also typical of tau filaments extracted from brains and spinal cords of individuals with Guam and Kii ALS/PDC, suggesting that similar molecular mechanisms underlie these diseases.

## RESULTS

### Structural characterisation of filaments from Guam ALS/PDC

We used cryo-EM to characterise filaments from the frontal cortex of 3 cases of Guam ALS/PDC and the spinal cord of cases 2 and 3 (Figure 1; Supplementary Figures 1-5). Staining with anti-tau antibody AT8 showed abundant neurofibrillary tangles (intracellular and extracellular) in frontal cortex (Supplementary Figure 6). As described (12), tau inclusions were also found in astrocytes and oligodendrocytes, with astrocytic inclusions mostly in subpial and perivascular areas.

**Figure 1:**
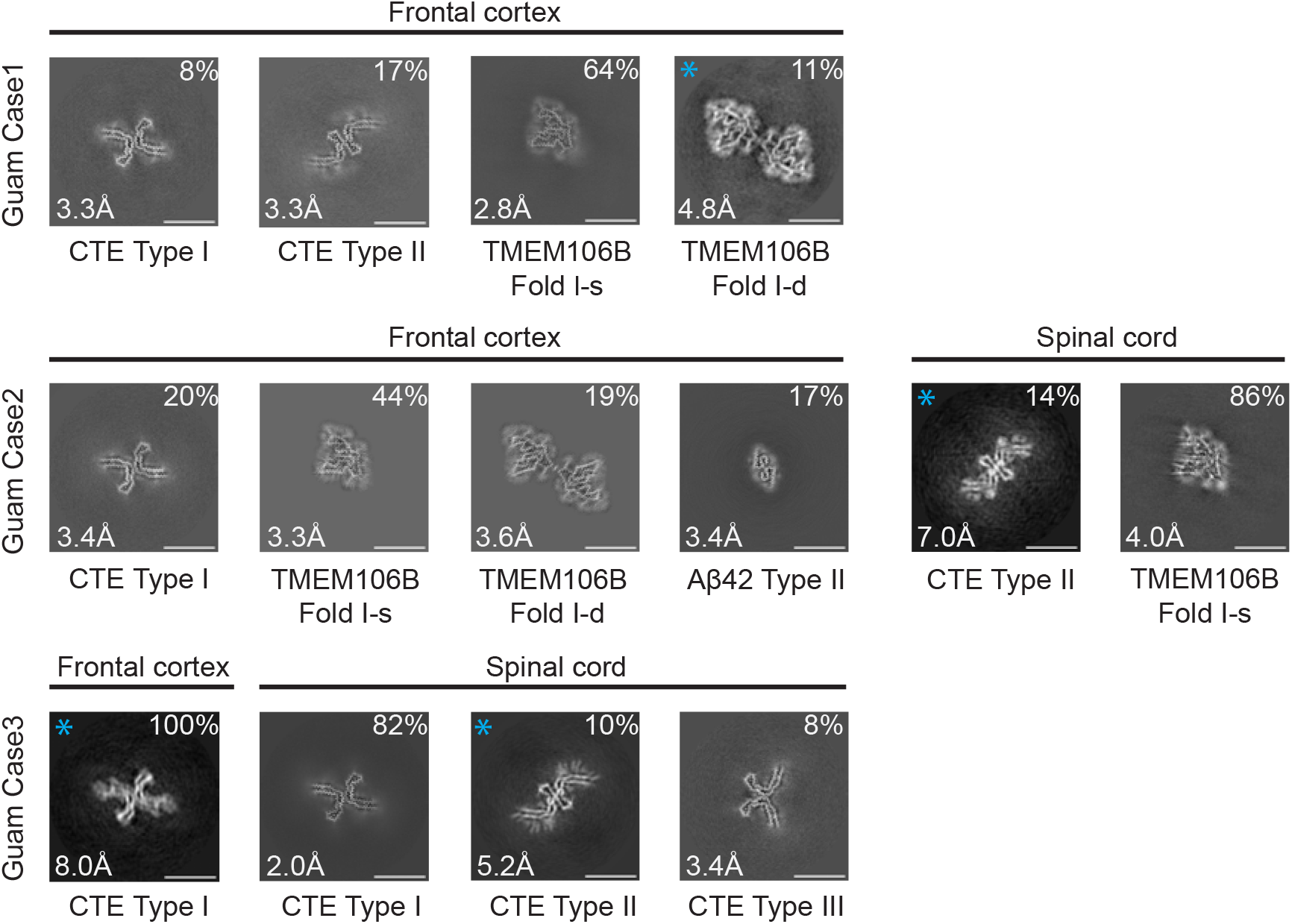
Cross-sections of cryo-EM structures of filaments from Guam ALS/PDC. Cross-sections perpendicular to the helical axis of cryo-EM structures of filaments from frontal cortex and spinal cord of 3 cases of Guam ALS/PDC, with a projected thickness of approximately one rung along the helical axis. For filament types indicated with an asterisk, there were insufficient images for high-resolution reconstruction and identification of the filament types was also based on 2D class averages (Supplementary Figure 1). Filament types are indicated, as are structural resolution and percentages of each filament type. Scale bar, 10 nm.

In all five samples, tau filaments with the CTE fold were present. The frontal cortex from case 1 contained a mixture of Type I and Type II filaments, whereas that from cases 2 and 3 had only Type I filaments. The spinal cord from case 2 had only Type II filaments, whereas that from case 3 contained a mixture of Type I and Type II filaments. In addition to tau filaments, we also observed singlets and doublets of transmembrane protein 106B (TMEM106B) filaments (fold I) in the frontal cortex from cases 1 and 2, and TMEM106B singlets (fold I) in the spinal cord from case 2 (Figure 1, Supplementary Figures 1 and 2). The frontal cortex from case 2 also contained Type II Aβ42 filaments (Figure 1, Supplementary Figures 1 and 2), like those that were described in brain extracts from cases of AD and other diseases (36). For several filament types, there were insufficient images for *de novo* three-dimensional reconstruction to high resolution. Their identification was also based on 2D class averages (Figure 1, Supplementary Figure 1).

High-resolution structure determination confirmed that the tau filament structures from Guam ALS/PDC are identical to those from CTE (Figure 2a). The root mean square deviation (RMSD) of Cα atoms in one rung of the filaments between Type I filaments from the spinal cord of Guam case 3 and those from CTE (PDB:6NWP) was 0.28 Å; the RMSD between Type II filaments from the frontal cortex of Guam case 1 and those from CTE (PDB:6NWQ) was 1.36 Å.

**Figure 2:**
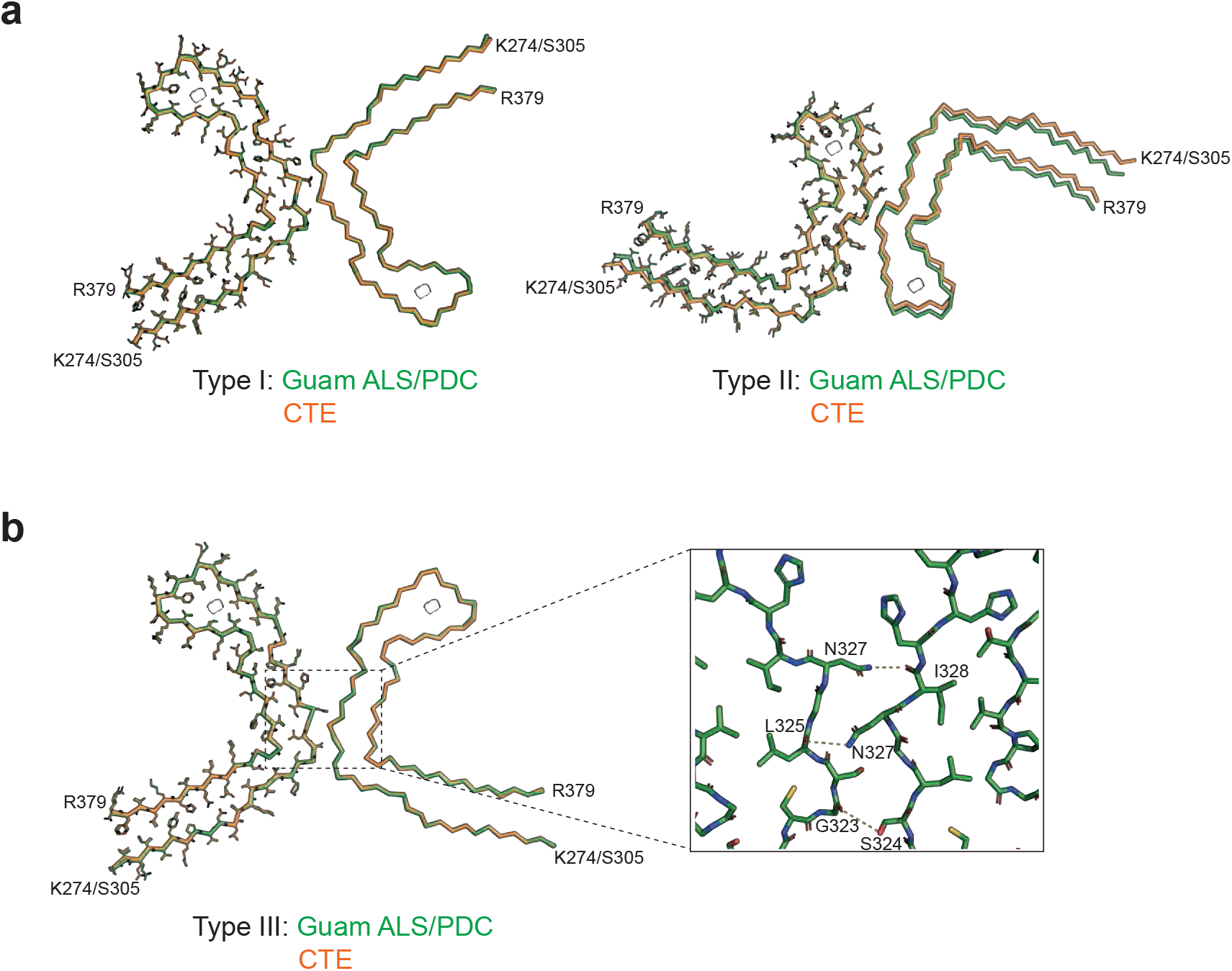
Comparison of tau filaments from Guam ALS/PDC and CTE. **(a)**, Overlay of the full atomic model (left protofilaments) and the main-chain trace (right protofilaments) of Guam ALS/PDC (green) and CTE Type I (orange) filaments (left) and Guam ALS/PDC (green) and Type II (orange) filaments (right). **(b)**, As in (a), but for Guam ALS/PDC Type III filaments (green) and CTE Type III filaments (orange). Inset: Zoomed-in view of the inter-protofilament packing of Guam ALS/PDC Type III filaments.

In the spinal cord of Guam case 3, we found a small proportion of filaments (less than 10%) with a previously unobserved structure, which we named CTE Type III tau filaments (Figures 1, 2b, Supplementary Figure 5). Two protofilaments with the CTE fold, spanning residues K274-R379 of three-repeat tau and S305-R379 of four-repeat tau, pack against each other in a back-to-back manner. The mirror-like arrangement of protofilaments in the XY cross-section indicates that they adopt opposite polarities in the filaments: one protofilament is oriented from top to bottom, while the other is oriented from bottom to top. The protofilament interface consists of residues ^323^GSLGNIH^329^ from both protofilaments, like in the CTE Type I filament interface. However, they form a different, staggered parallel zipper, in which the side chains of S324 and N327 of both protofilaments intercalate and form hydrogen bonds with the main chain groups of opposite protofilaments (Figure 2b). As in Type I and Type II filaments, both protofilaments in Type III filaments harbor an additional density in the β-helix region (Figure 2b, Supplementary Figures 2 and 5). CTE Type III tau filaments were also found in new cryo-EM images of filaments from the temporal cortex of an individual with CTE [case 2 in (30)] (Supplementary Figure 5), indicating that they are not restricted to Guam ALS/PDC. The relatively small numbers of filaments with the CTE Type III fold probably explain why this type of filament had not been detected previously.

### Structural characterisation of filaments from Kii ALS/PDC

We analysed extracts from temporal cortex of 8 cases of ALS/PDC from the Kii peninsula (Figure 3, Supplementary Figures 1-4). Staining with AT8 showed the presence of abundant neurofibrillary tangles that were particularly abundant in cortical layers II/III (Supplementary Figure 7). Tau-positive astrocytes and coiled bodies were also present. Case 8 has previously been shown to exhibit astrocytic plaque-like structures and threads, reminiscent of corticobasal degeneration (11).

**Figure 3:**
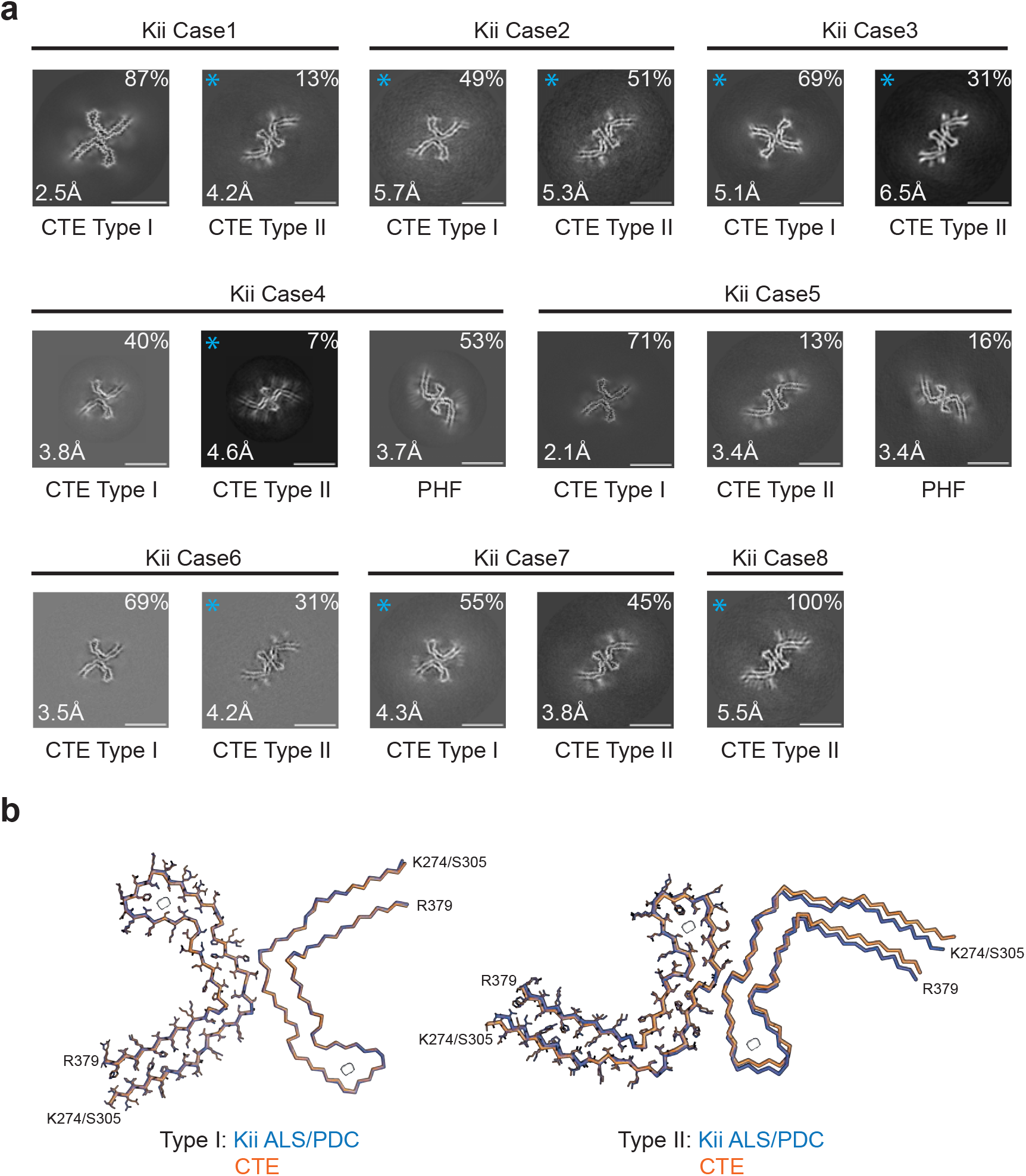
Cross-sections of cryo-EM structures of filaments from Kii ALS/PDC and comparison of tau filaments with those from CTE. **(a)**, Cross-sections perpendicular to the helical axis of cryo-EM structures of filaments from 8 cases of Kii ALS/PDC, with a projected thickness of approximately one rung along the helical axis. For filament types indicated with an asterisk, there were insufficient images for high-resolution reconstruction and identification of the filament types was also based on 2D class averages (Supplementary Figure 1). Filament types are indicated, as are structural resolution and percentages of each filament type. Scale bar, 10 nm. **(b)**, Overlay of the full atomic model (left protofilaments) and the main-chain trace (right protofilaments) of Kii ALS/PDC (blue) and CTE Type I (orange) filaments (left) and Kii ALS/PDC (green) and Type II (orange) filaments (right).

Again, all the samples contained tau filaments with the CTE fold. Case 6 had only Type II filaments; all other cases had a mixture of Type I and Type II filaments. Cases 2 and 5 also contained tau paired helical filaments (PHFs), like those from AD and other conditions (28,29,31,32). We did not observe Aβ or TMEM106B filaments. High-resolution structure determination showed that the structures of tau filaments from Kii ALS/PDC are also the same as those from CTE (Figure 3b). The RMSD between Type I filaments from Kii case 2 and those from CTE (PDB:6NWP) was 0.38 Å; the RMSD between Type II filaments from Kii case 2 and those from CTE (PDB:6NWQ) was 1.37 Å.

## DISCUSSION

Abundant filamentous amyloid inclusions that are made of all six brain tau isoforms are characteristic of ALS/PDC (8,13). We now show that tau filaments from Guam and Kii ALS/PDC adopt the CTE fold (30) in brain and spinal cord. We recently showed that tau filaments from SSPE also adopt the CTE fold (33).

These findings suggest that the molecular mechanisms that cause tau assembly in ALS/PDC may be similar to those at work in CTE and SSPE. The latter two are probably caused by environmental factors, in the form of repetitive head injuries and measles infection, respectively. Neuroinflammation may be important in both diseases. Exogenous factors may also be causal in Guam and Kii ALS/PDC, with a possible role for parasitic infestation (21,22).

As in CTE (26) and SSPE (35), more filamentous tau inclusions in ALS/PDC of Guam and Kii are found in layers II/III of the cerebral cortex than in layers V/VI (10,11). This is unlike AD, where tau inclusions are more abundant in layers V/VI (25). The presence of Alzheimer and CTE tau folds correlates with these differences. It suggests that the CTE fold may also form in other diseases with a predominance of tau inclusions in cortical layers II/III that are believed to be caused by environmental factors, such as postencephalitic parkinsonism (37) and the Nodding syndrome (38).

The CTE tau fold differs from the Alzheimer fold by having a more open conformation of the β-helix region, which contains an internal density of unknown identity (30). In the presence of NaCl, recombinant tau comprising residues 297-391 assembled into filaments with the CTE fold, but in its absence, the Alzheimer tau fold formed (39). It remains to be seen how this difference relates to human brains.

Besides tau filaments with the CTE fold, we also observed Type II Aβ42 filaments in a case from Guam and tau PHFs in two cases from the Kii peninsula. Senile plaques have been described in around 60% of cases of Guam ALS/PDC (15) and assembly of Aβ is believed to be part of the disease process (40). Alternatively, these changes may be age-related. This was probably also the reason for the presence of TMEM106B filaments (41,42) in two cases from Guam. It is possible that Aβ and TMEM106B filaments were lost during the extraction method used for the Kii cases. In addition to tau, also Aβ, α-synuclein and TDP-43 inclusions have been implicated in the pathogenesis of ALS/PDC (11,14,15). We did not find α-synuclein or TDP-43 filaments.

In conclusion, we demonstrate the presence of tau filaments with the CTE fold in cases of ALS/PDC from the island of Guam and the Kii peninsula. Type I and/or Type II CTE filaments were present in brains and spinal cords. We also describe the new CTE Type III tau filament, in which two protofilaments pack with opposite polarities. The presence of tau filaments with the CTE fold supports the hypothesis that ALS/PDC is caused by exogenous factors.

## MATERIALS AND METHODS

### Cases of ALS/PDC

Three cases of ALS/PDC from the island of Guam and 8 cases from the Kii peninsula were investigated. The Guam cases have not been reported before; we used tissues from two Chamorro males and one ½ Chamorro, ½ Filippina female with long-standing dementia and Parkinson’s disease, in the absence of a family history of disease. They belonged to the PDC subtype, where some tau inclusions can be found in spinal cord (43). They died aged 73 (cases 1 and 2) and aged 69 (case 3). The cases from the Kii peninsula have been published (11). Three individuals (cases 1,2,5) belonged to the ALS subtype and 5 (cases 3,4,6-8) to the PDC subtype. The ages at death were: ALS subtype, 63, 76 and 77 years; PDC subtype, 60, 70, 71, 74 and 74 years. The duration of illness varied between 1-14 years. There was no history of head injury or measles infection in either the Guam or the Kii cases of ALS/PDC. This study was approved by the Ethics Committees of the Universities of Shinshu (3233 and 5108), Niigata (2020-0019) and Mie (2592).

### Immunohistochemistry

Brains were fixed in 20% buffered formalin, cut into coronal sections and paraffin-embedded. Sections (4.5 μm) were incubated overnight at room temperature with antibody AT8, which is specific for pS202 and pT205 tau (1:5,000, Innogenetics) (44). To reveal the signal, the Envision plus kit (Dako) was used, with diaminobenzidine tetrahydrochloride (Sigma-Aldrich) as chromogen. Some sections from Kii cases of ALS/PDC were also stained with Gallyas-Braak silver (45).

### Filament extraction

For the Guam ALS-PDC cases, sarkosyl-insoluble material was extracted from frontal cortex (cases 1-3) and spinal cord (cases 2 and 3), as described (46). The tissues (less than 100 mg) were homogenised in 3ml buffer A (10 mM Tris-HCl, pH 7.5, 0.8 M NaCl, 10% sucrose and 1 mM EGTA), brought to 2% sarkosyl and incubated for 30 min at 37° C. The samples were centrifuged at 7,000g for 10 min, followed by spinning the supernatants at 100,000g for 60 min. The pellets were resuspended in 100 μl/g of buffer B (20 mM Tris-HCl, pH 7.4, 100 mM NaCl) for cryo-EM analysis. For ALS/PDC cases from the Kii peninsula, minor changes were made to the above extraction protocol. After incubation in 2% sarkosyl, the samples were sonicated (TAITEC ultrasonic homogeniser VP-55, level 7) for 15 s and, following a 10 min centrifugation at 27,000g, supernatants were centrifuged at 257,400g for 30 min at 25° C. The pellets were then resuspended in 900 μl/g buffer A with 1% sarkosyl and centrifuged at 166,000g for 20 min at 25° C. Filaments from the CTE brain [case 2 in (30)] were extracted as described (47).

### Electron cryo-microscopy

Three μl of the sarkosyl-insoluble fractions were applied to glow-discharged (Edwards S150B) holey carbon grids (Quantifoil Au R1.2/1.3, 300 mesh) that were plunge-frozen in liquid ethane using a Vitrobot Mark IV (Thermo Fisher Scientific) at 100% humidity and 4°C. Cryo-EM images were collected on a Titan Krios electron microscope operated at 300 kV and equipped with a Falcon-4 or a K3 direct electron detector. Images were recorded in electron event representation (EER) format (48) for Falcon-4 (6s) and Tif format for K3 (1s), with a total dose of 40e/Å^2^ and a pixel size of 0.824 Å (Falcon-4) or 0.826 Å (K3). See Supplementary Tables 1 and 2 for further details.

### Helical reconstruction

Datasets were processed in RELION using standard helical reconstruction (49). Movie frames were gain corrected, aligned and dose weighted using RELION’s own motion correction program (50). Contrast transfer function (CTF) parameters were estimated using CTFFIND4-1 (51). Filaments were picked manually. For the analysis of filament types and the generation of initial three-dimensional models, segments were extracted with a box size of 1024 pixels and down-scaled to 256 pixels. Reference-free 2D classification was performed to discard suboptimal images and to measure cross-over distances for initial model calculation using relion_helix_inimodel2d (52). For high-resolution refinement, selected segments were extracted with a box size of 400 pixels, with the original pixel size. 3D auto-refinements were performed with optimisation of the helical twist and rise parameters once resolutions extended beyond 4.7 Å. To improve the resolution, Bayesian polishing and CTF refinement were performed (53). Final maps were sharpened using standard post-processing procedures in RELION and resolution estimates calculated based on the Fourier shell correlation (FSC) between two independently refined half-maps at 0.143 (54).

### Model building and refinement

Atomic models were built manually in Coot (55), based on published structures [CTE type I, PDB:6NWP; CTE type II, PDB:6NWQ; TMEM106B fold I-s, PDB:7QVC; TMEM106B fold I-d, PDB:7QVF; Type II Aβ42, PDB:7Q4M (30,36,41). Model refinements were performed using *Servalcat* (56) and REFMAC5 (57,58). Models were validated with MolProbity (59). Figures were prepared with ChimeraX (60) and Pymol (61).

## ACKNOWLEDGEMENTS

We are grateful to the patients’ families for donating brain tissues. We thank members of the EM facility at the MRC Laboratory of Molecular Biology for support with data acquisition and Jake Grimmett, Toby Darling and Ivan Clayson for help with high-performance computing. This work was funded by the Medical Research Council, as part of U.K. Research and Innovation (MC_UP_120/25 to B.R.-F., MC_UP_A025_1013 to S.H.W.S. and MC_U105184291 to M.G.). It was also supported by the U.S. National Institute on Aging (R01AG054641, to M.V. and B.M.V.), a Pilot Award from the Southern California Environmental Health Sciences Center (to M.V. and B.M.V.), the Mie Medical Fund (to Y.K. and S.M.), the Research Committee of CNS Degenerative Diseases (to Y.K.), the Research Committee on Muro Disease (21210301 to Y.K.), the Japan Society for the Promotion of Science (JSPS KAKENHI, JP18K07514 to Y.K. and JP22H04923 to S.M.), the Japan Agency for Medical Research and Development (to Y.K., M.Y. and M.H.), a Swiss National Science Postdoctoral Fellowship (P500PB 206890 to S.T.) and a Collaborative Research Project of the Niigata Brain Research Institute (to A.K.). We acknowledge the contributions of Professor Shigeki Kuzuhara (1944-2021) to the study of Kii ALS/PDC. For the purpose of open access, the MRC Laboratory of Molecular Biology has applied a CC-BY public copyright licence to any Author Accepted Manuscript version arising.

## AUTHOR CONTRIBUTIONS

B.M.V., Y.K., A.N., S.M., M.V., R.S., E.A., Y.H., K.O., A.K. and M.Y. identified the patients and performed neuropathology; C.Q., Y.S., S.T. and M.H. extracted filaments; C.Q., Y.S. and S.T. acquired cryo-EM data; C.Q., Y.S., S.T., A.G.M., B.R.-F. and S.H.W.S. did structure determination; S.H.W.S. and M.G. supervised the project and all authors contributed to the writing of the manuscript.

## DATA AVAILABILITY

Cryo-EM maps have been deposited in the Electron Microscopy Data Bank (EMDB) with the following accession numbers: EMD-17171, EMD-17173, EMD-17174, EMD-17175, EMD-17176, EMD-17177, EMD-17178, EMD-17179, EMD-17180, EMD-17181. Corresponding refined atomic models have been deposited in the Protein Data Bank (PDB) under the following accession numbers: 8OT6, 8OTC, 8OTD, 8OTE, 8OTF, 8OTG, 8OTH, 8OTJ, 8OTI. Please address requests for materials to the corresponding authors.

## SUPPLEMENTARY FIGURES

**Supplementary Figure 1:**
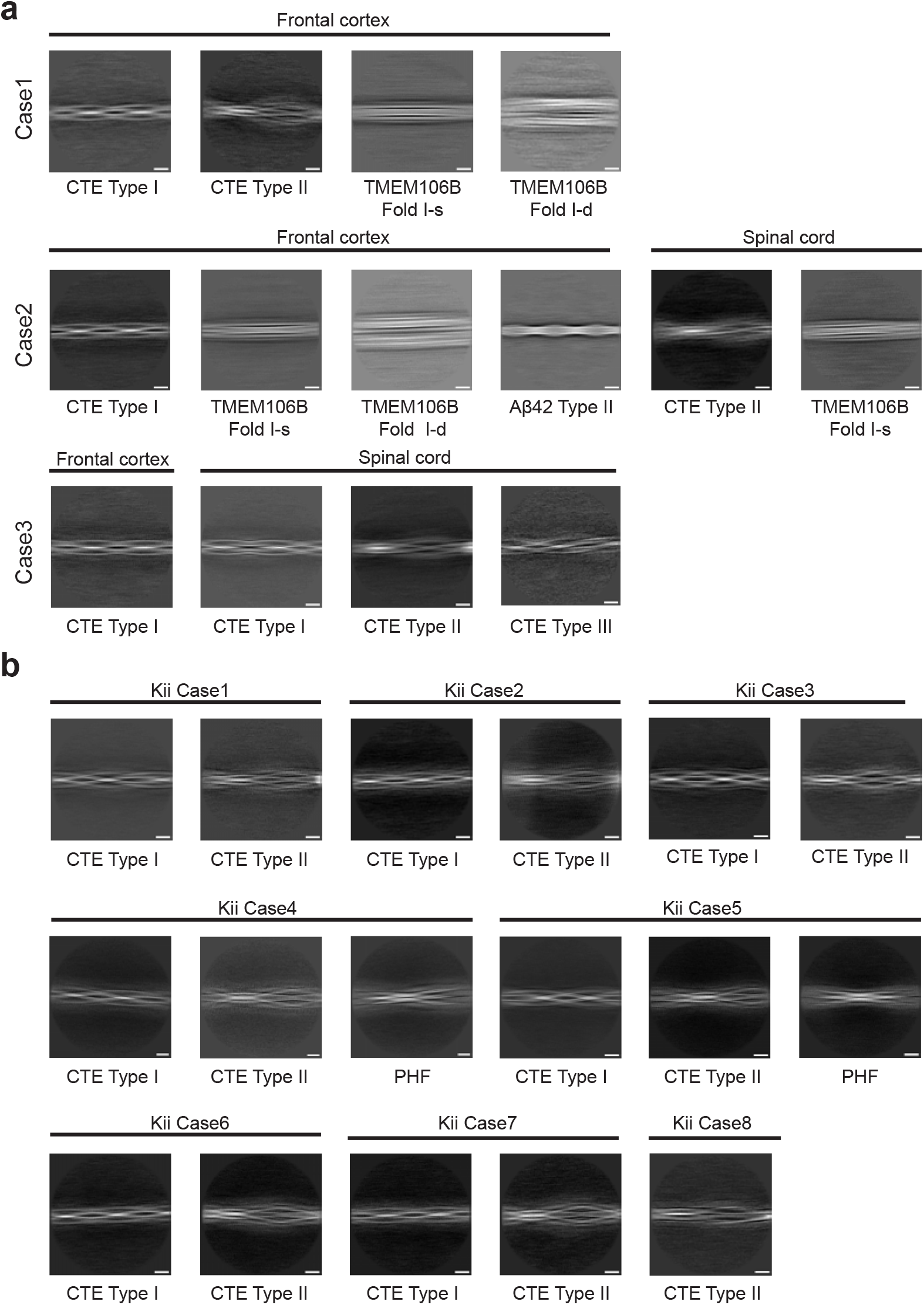
Two-dimensional classification of filaments from Guam (a) and Kii (b) ALS/PDC. Scale bar, 10 nm.

**Supplementary Figure 2:**
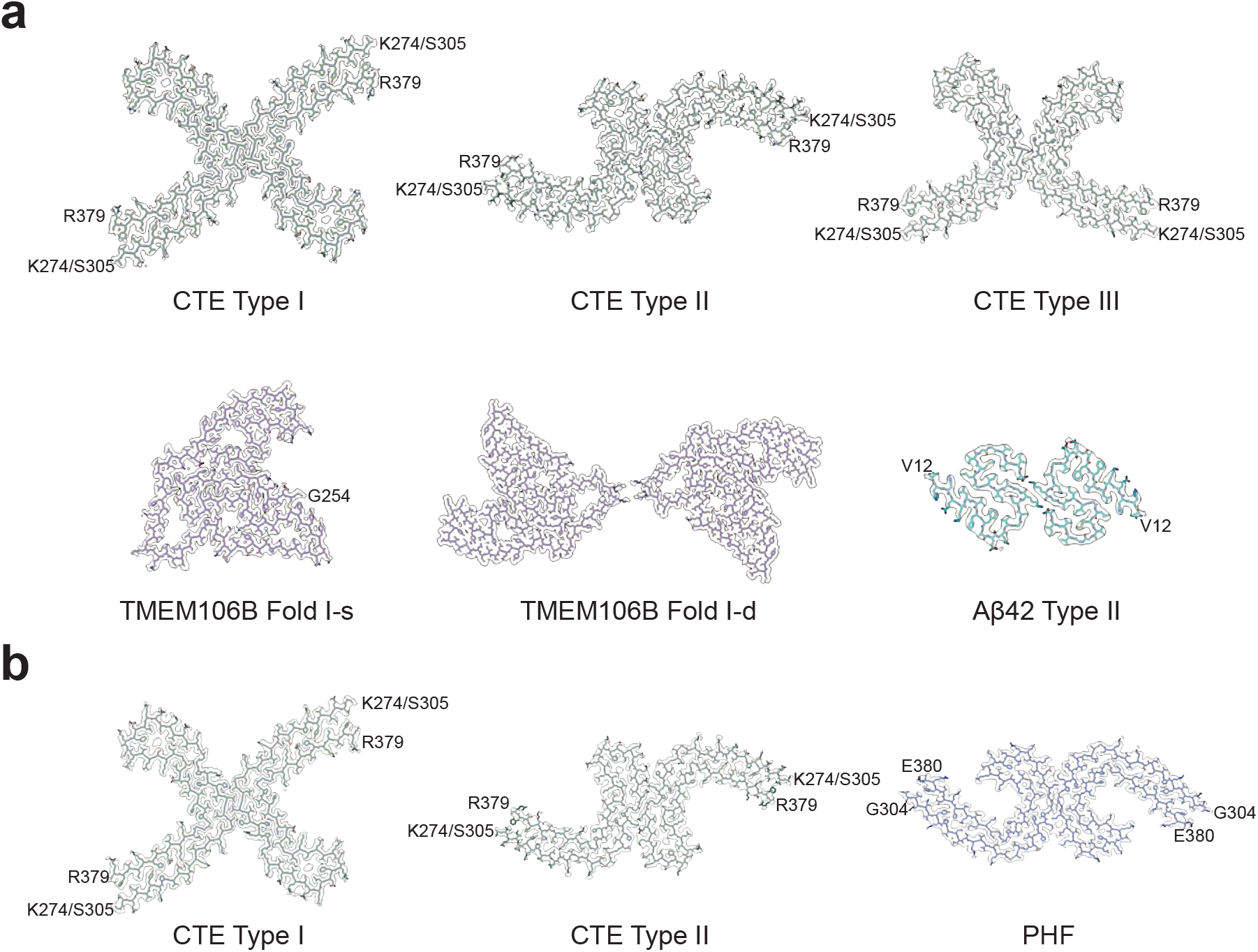
Cryo-EM density maps of filaments from Guam and Kii ALS/PDC. **(a)**, In Guam ALS/PDC, Type I, Type II and Type III tau filaments (green), singlets and doublets of TMEM106B filaments (fold I) (purple) and Type II Aβ42 filaments (cyan) were present. **(b)**, In Kii ALS/PDC, Type I and Type II tau filaments (green), as well as tau paired helical filaments (PHFs) (blue), were present.

**Supplementary Figure 3:**
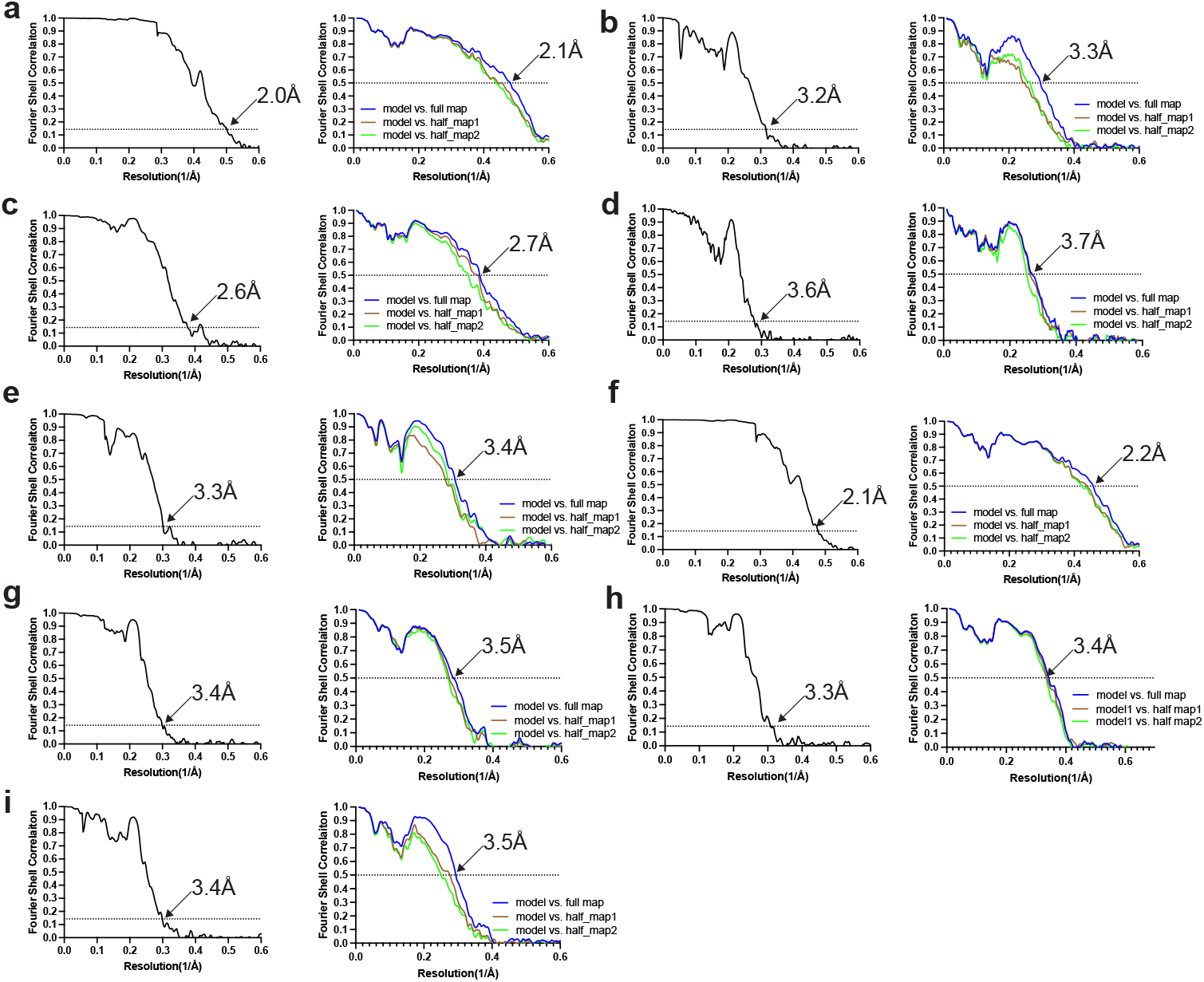
Fourier shell correlation (FSC) curves. FSC curves of cryo-EM maps (left panel) and model to map validation (right panel). **(a)**, Guam ALS/PDC CTE tau Type I. **(b)**, Guam ALS/PDC CTE tau Type II. **(c)**, Guam ALS/PDC TMEM106B fold I-s. **(d)**, Guam ALS/PDC TMEM106B fold I-d. **(e)**, Guam ALS/PDC CTE tau Type II Aβ42. **(f)**, Kii ALS/PDC CTE tau Type I. **(g)**, Kii ALS/PDC CTE tau Type II. **(h)**, Kii ALS/PDC tau PHF. **(i)**, Guam ALS/PDC CTE tau Type III.

**Supplementary Figure 4:**
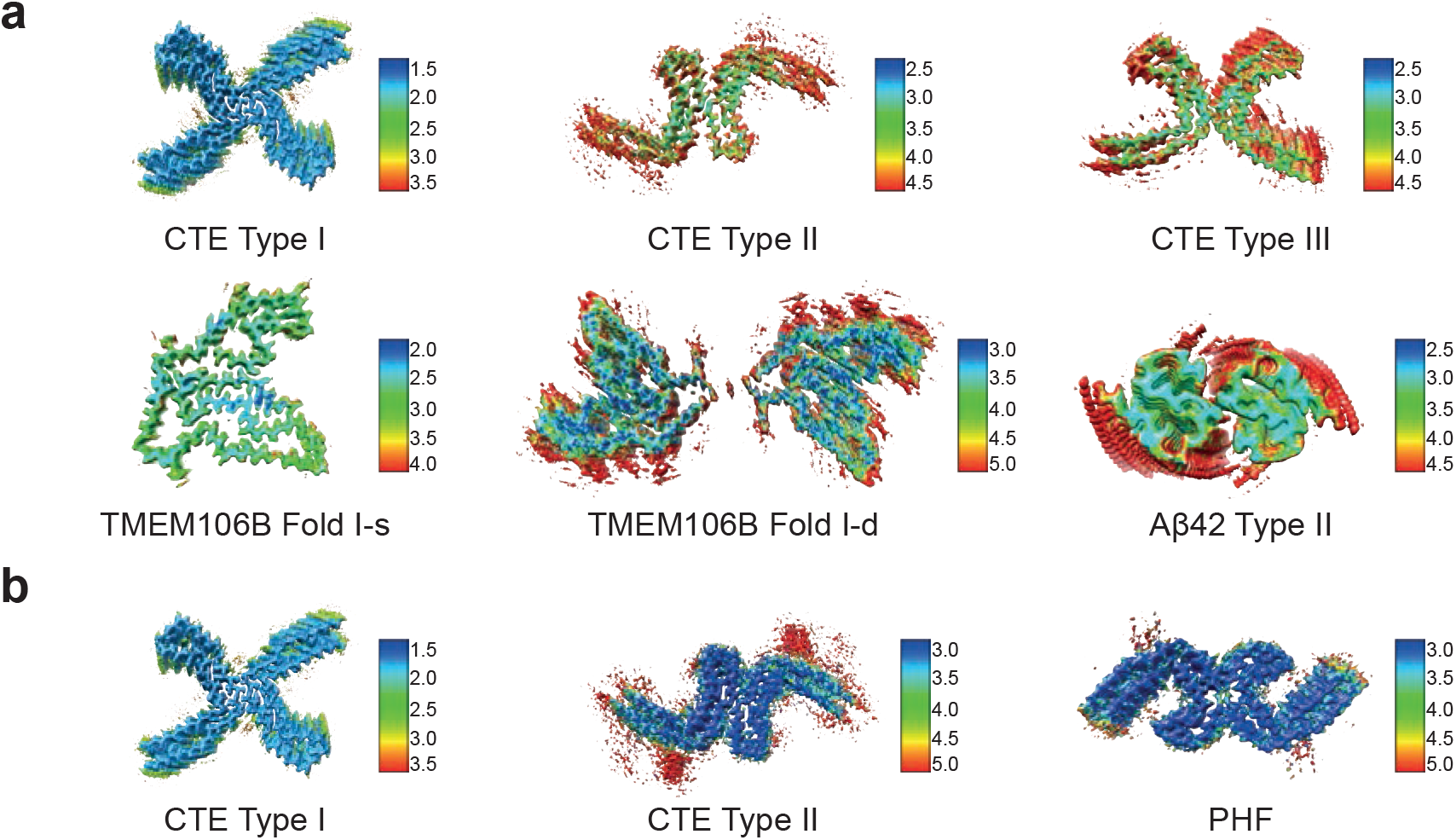
Local resolution estimation of filaments from Guam (a) and Kii (b) ALS/PDC.

**Supplementary Figure 5:**
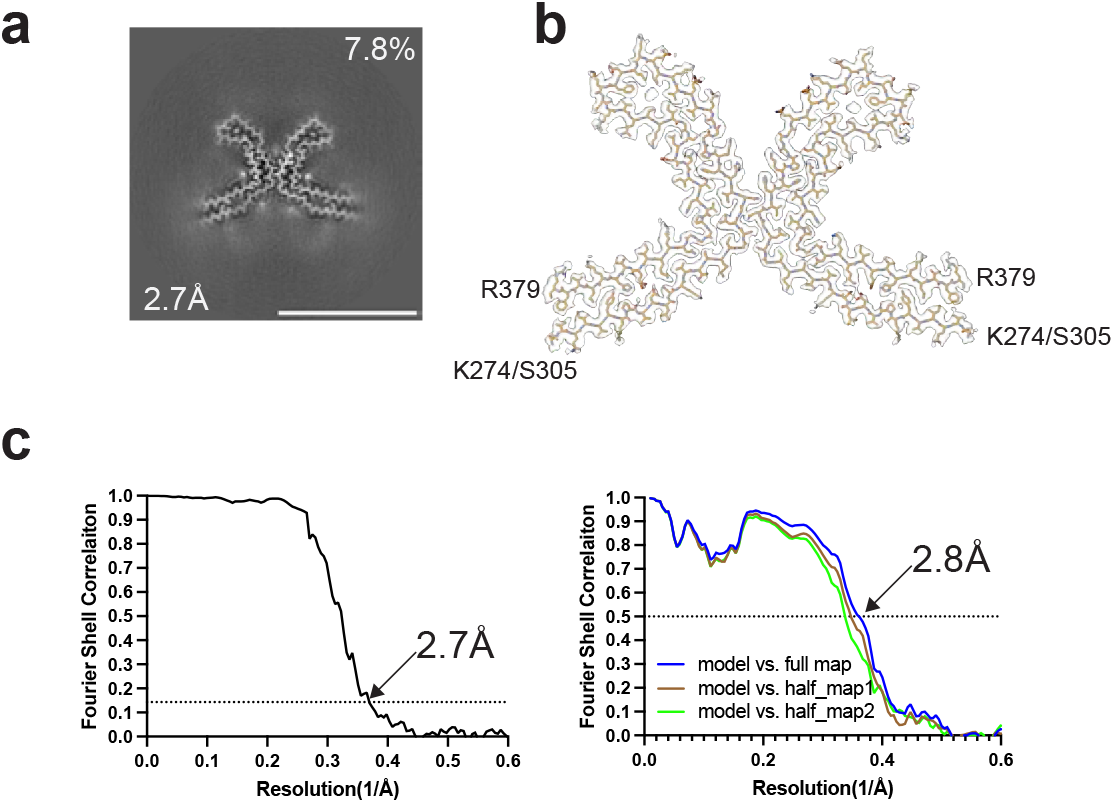
Type III filaments from CTE. **(a)**, Cross-section perpendicular to the helical axis of the cryo-EM structure of Type III filaments from temporal cortex of CTE case 2 (30), with a projected thickness of approximately one rung along the helical axis. **(b)**, Cryo-EM density map and model of Type III filament. **(c)**, Fourier shell correlation curves of cryo-EM maps of Type III filaments (left panel) and model to map validation (right panel).

**Supplementary Figure 6:**
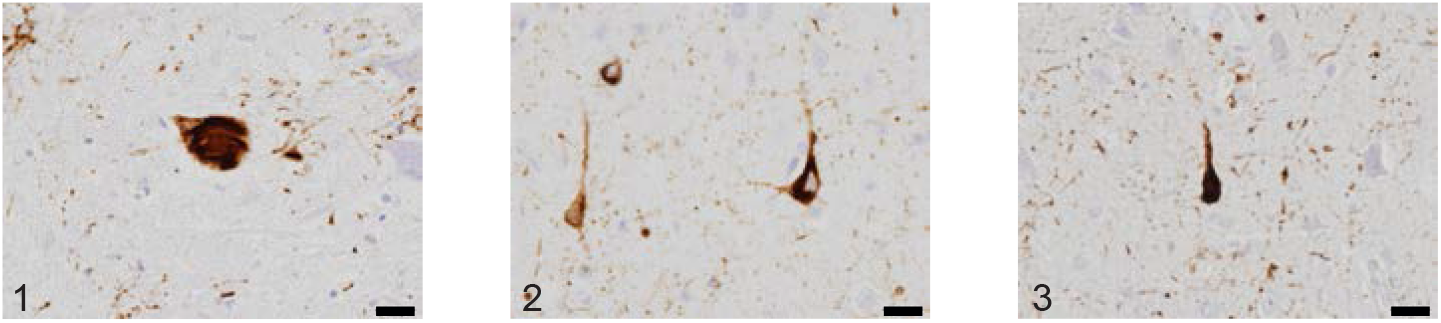
Immunostaining of tau inclusions from Guam ALS/PDC. Sections from the frontal cortex of cases 1-3 stained with anti-tau antibody AT8. Scale bar, 20 μm.

**Supplementary Figure 7:**
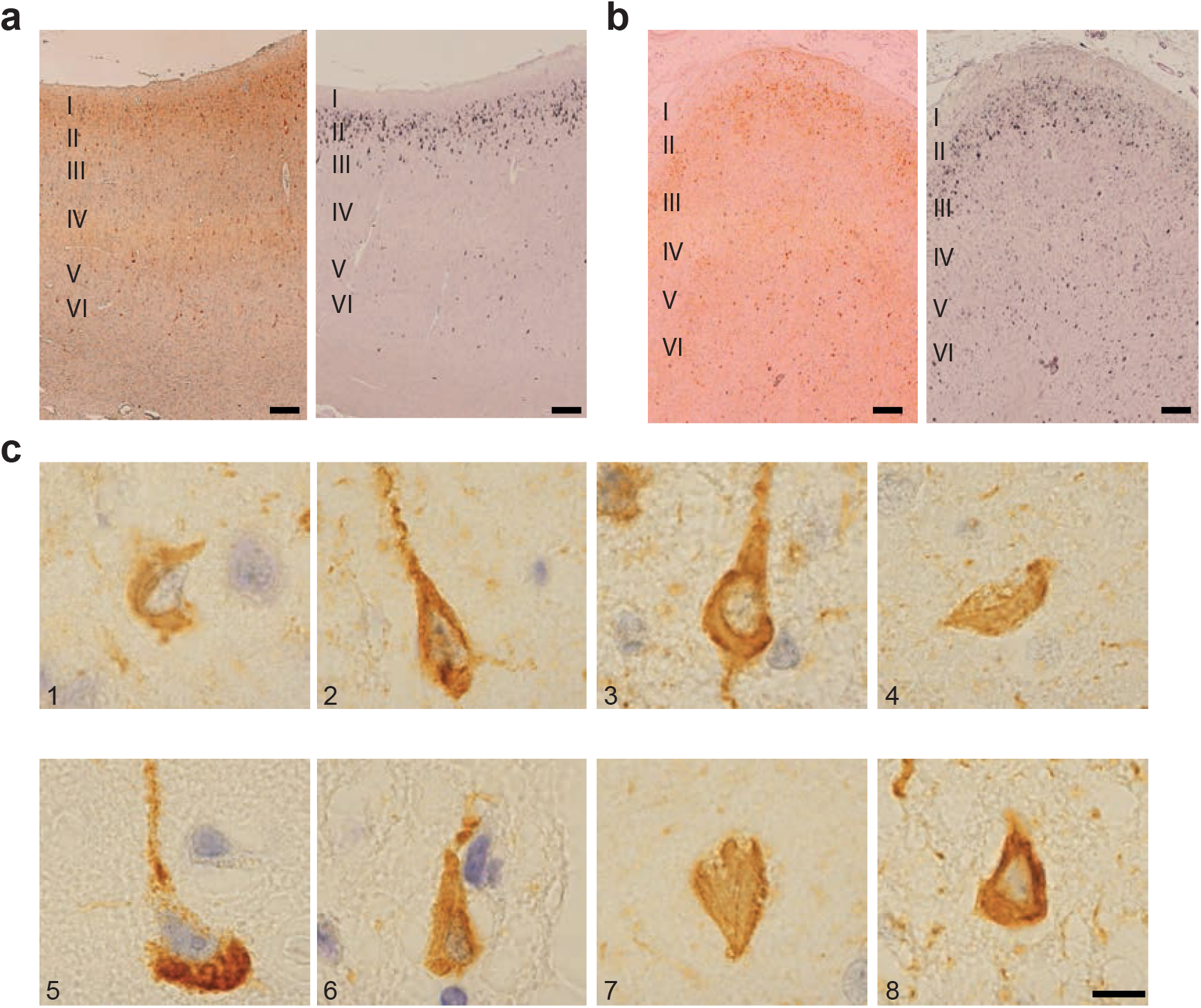
Immunostaining of tau inclusions from Kii ALS/PDC. **(a,b)**, Temporal cortex from cases 3 and 4 stained with anti-tau antibody AT8 (left) and with Gallyas-Braak silver (right). Tau inclusions are concentrated in layers II/III. Scale bar, 200 μm. **(c)**, Sections from the temporal cortex of cases 1-8 stained with AT8. Scale bar, 10 μm.

## SUPPLEMENTARY TABLES

**Supplementary Table 1:** Cryo-EM data collection, refinement and validation statistics for Guam ALS/PDC.

|  | Guam Case3 spinal cord |  | Guam Case1 frontal cortex |  | Guam Case2 frontal cortex |  |
| --- | --- | --- | --- | --- | --- | --- |
| **Data collection** |  |  |  |  |  |  |
| Microscope | Titan Krios |  | Titan Krios |  | Titan Krios |  |
| Voltage (kV) | 300 |  | 300 |  | 300 |  |
| Detector | Falcon4 |  | Falcon4 |  | Falcon4 |  |
| Magnification | 96,000 |  | 96,000 |  | 96,000 |  |
| Electron exposure (e-/Å<sup>2</sup>) | 40 |  | 40 |  | 40 |  |
| Defocus range (μm) | -1.0 to -2.0 |  | -1.0 to -2.0 |  | -1.0 to -2.0 |  |
| Pixel size (Å) | 0.824 |  | 0.824 |  | 0.824 |  |
| **Data processing** | CTE TypeI | CTE TypeIII | CTE type II | TMEM106B<br>Fold I-s | TMEM106B<br>Fold I-d | Aβ42 TypeII |
| Box size (pixel) | 400 | 400 | 400 | 400 | 400 | 400 |
| Symmetry imposed | C1 | C1 | C1 | C1 | C1 | C2 |
| Initial particle images (no.) | 256,999 |  | 130,240 |  | 281,921 |  |
| Final particle images (no.) | 140,124 | 13,456 | 3,105 | 57,802 | 11,131 | 20,363 |
| Map resolution (Å)<br>FSC threshold 0.143 | 2.0 | 3.4 | 3.2 | 2.6 | 3.6 | 3.3 |
| Helical rise (Å) | 2.37 | 4.74 | 2.38 | 4.81 | 4.79 | 4.79 |
| Helical twist (°) | 179.41 | -1.19 | 179.4 | -0.4 | -0.42 | -2.99 |
| **Refinement** |  |  |  |  |  |  |
| Model resolution (Å)<br>FSC threshold 0.5 | 2.1 | 3.5 | 3.3 | 2.7 | 3.7 | 3.4 |
| Map sharpening *B* factor (Å<sup>2</sup>) | -26 | -54 | -30 | -28 | -69 | -43 |
| Model composition |  |  |  |  |  |  |
| Non-hydrogen atoms | 2870 | 3444 | 3444 | 4340 | 8680 | 1356 |
| Protein residues | 375 | 450 | 450 | 540 | 1080 | 186 |
| Ligands | 0 | 0 | 0 | 0 | 0 | 0 |
| *B* factors (Å<sup>2</sup>) |  |  |  |  |  |  |
| Protein | 104.2 | 196.3 | 312.4 | 141.1 | 222.2 | 156.8 |
| R.m.s. deviations |  |  |  |  |  |  |
| Bond lengths (Å) | 0.004 | 0.007 | 0.0056 | 0.0056 | 0.0071 | 0.0073 |
| Bond angles (°) | 1.006 | 1.541 | 1.11 | 1.367 | 1.537 | 1.369 |
| Validation |  |  |  |  |  |  |
| MolProbity score | 0.95 | 2.06 | 1.39 | 2.21 | 2.54 | 2.17 |
| Clashscore | 0.51 | 3.56 | 1.99 | 4.15 | 5.19 | 4.71 |
| Poor rotamers (%) | 1.52 | 6.82 | 1.52 | 4.76 | 5.95 | 4.35 |
| Ramachandran plot |  |  |  |  |  |  |
| Favored (%) | 97.26 | 95.89 | 95.89 | 91.73 | 84.21 | 93.1 |
| Allowed (%) | 2.74 | 4.11 | 4.11 | 8.27 | 15.79 | 6.9 |
| Disallowed (%) | 0 | 0 | 0 | 0 | 0 | 0 |
| PDB | 8OT6 | 8OT9 | 8OTC | 8OTD | 8OTE | 8OTF |
| EMDB | EMD-17171 | EMD-17173 | EMD-17174 | EMD-17175 | EMD-17176 | EMD-17177 |

**Supplementary Table 2:**
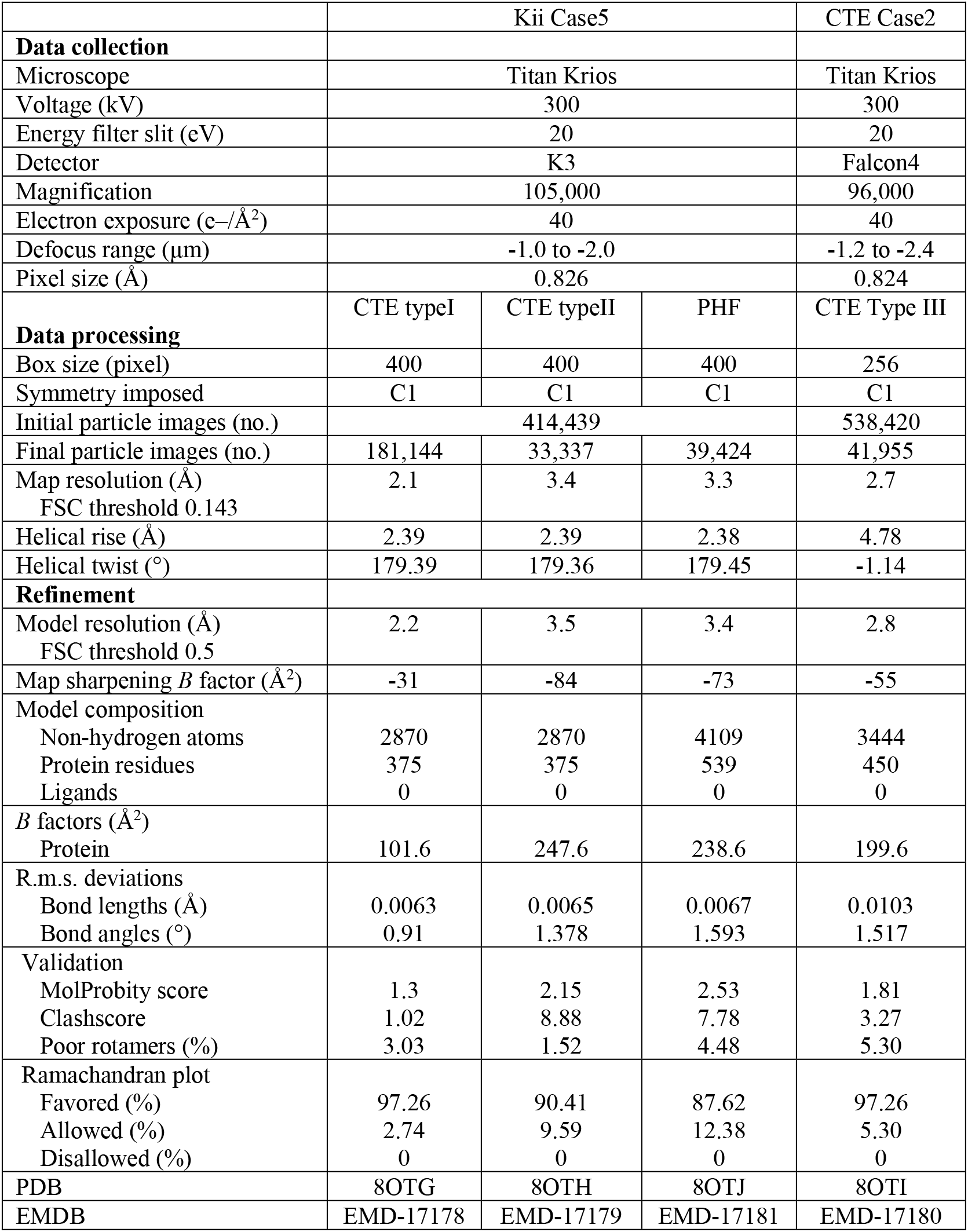
Cryo-EM data collection, refinement and validation statistics for Kii ALS/PDC and CTE.

